# Not so dry after all – DRY mutants of the AT1_A_ receptor and H1 receptor can induce G protein-dependent signaling

**DOI:** 10.1101/773044

**Authors:** A Pietraszewska-Bogiel, L Joosen, J Goedhart

## Abstract

GPCRs are seven transmembrane spanning receptors that regulate a wide array of intracellular signaling cascades in response to various stimuli. To do so, they couple to different heterotrimeric G proteins and adaptor proteins, including arrestins. Importantly, arrestins were shown to regulate GPCR signaling through G proteins, as well as promote G protein-independent signaling events. Several research groups have reported successful isolation of exclusively G protein-dependent and arrestin-dependent signaling downstream of GPCR activation using biased agonists or receptor mutants incapable of coupling to either arrestins or G proteins. In the latter category, the DRY mutant of the angiotensin II type 1 receptor was extensively used to characterize functional selectivity downstream of AT1_A_R. In an attempt to understand histamine 1 receptor signaling, we characterized the signaling capacity of the H1R DRY mutant in a panel of dynamic, live cell biosensor assays, including arrestin recruitment, heterotrimeric G-protein activation, Ca^2+^ signaling, protein kinase C activity, GTP binding of RhoA, and activation of ERK1/2. Here we show that both H1R DRY mutant and the AT1_A_R DRY mutant (used as a reference) are capable of efficient activation of G protein-mediated signaling. Therefore, contrary to common belief, they do not constitute suitable tools for dissection of arrestin-mediated, G protein-independent signaling downstream of these receptors.

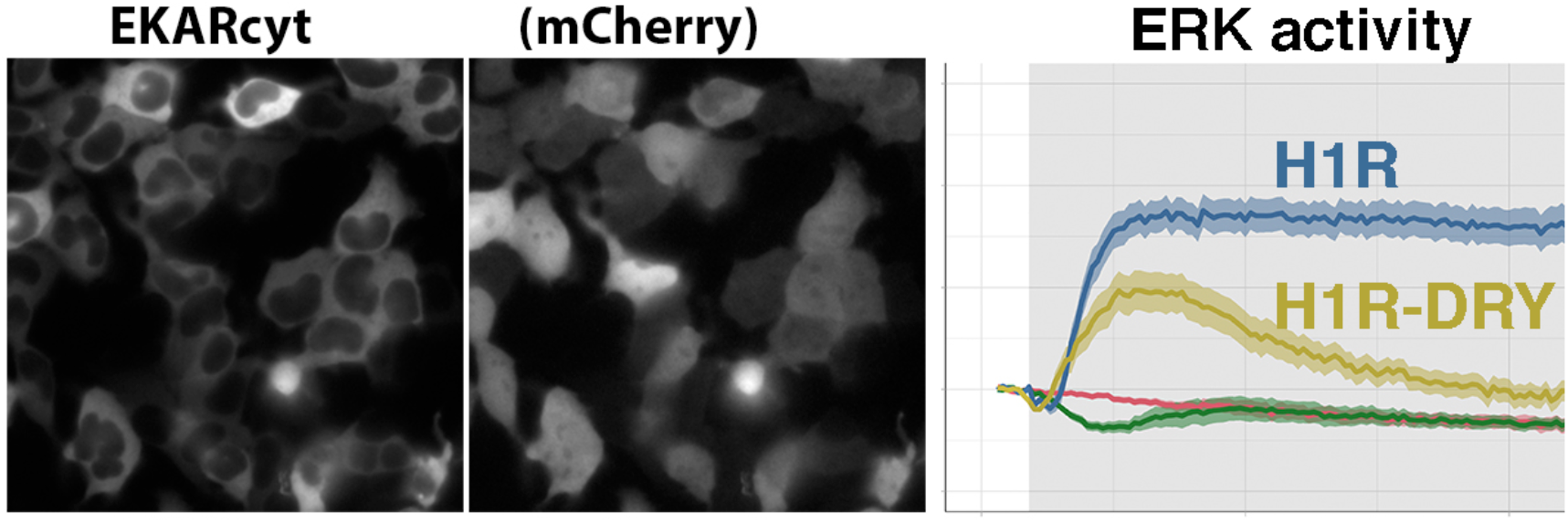

## INTRODUCTION

G protein-coupled receptors (GPCRs) constitute the largest family of cell surface proteins involved in signaling in response to various stimuli that underlie many cellular and physiological processes. Agonist binding to GPCRs evokes rearrangements in the intramolecular interactions within the seven transmembrane domain structure of the receptor that result in receptor activation and its coupling to heterotrimeric (Gαßγ) G proteins (reviewed in Weis & Kobilka 2018). This leads to G protein activation and switching of the canonical GPCR signaling via second messengers. Subsequent cessation of this G protein-dependent signaling occurs via recruitment of arrestins to the cytoplasmic surface of the receptor, a process that is enhanced by receptor phosphorylation by G protein-coupled receptor kinases, GRKs (reviewed in Hilger et al. 2018 and Komolov et al. 2018). Out of four arrestin isoforms, arrestin 1 and 4 bind photoreceptors in the retina, whereas two non-visual arrestins (arrestin 2 and 3 or ß-arrestin1 and 2, respectively) bind virtually all other GPCRs. Arrestin binding physically prevents receptor-G protein interaction, leading to desensitization of receptor-mediated activation of G proteins, and promotes the subsequent receptor endocytosis via clathrin-coated vesicles (Zhang et al. 1999, reviewed in Kang et al. 2014 and Gurevich & Gurevich 2015).

In addition to desensitization, arrestin binding to the receptor was proposed to “switch” the receptor from G protein signaling mode that transmits transient signals from the plasma membrane to arrestin signaling mode that transmits a distinct set of signals as the receptor internalizes (reviewed in Reiter et al. 2012 and Peterson & Luttrell 2018). Arrestin-mediated signaling downstream of GPCR activation is arguably most thoroughly studied in case of mitogen-activated, extracellular signal-regulated kinase 1 and 2 (ERK1/2) cascade. In fact, several research groups have reported successful “isolation” of G protein-independent ß-arrestin-mediated signaling downstream of GPCR activation using so called ß-arrestin-biased agonists (that do not support receptor coupling to G proteins; reviewed in Whalen et al. 2011, see also Saulière et al. 2012, Zimmerman et al. 2012, Littmann et al. 2015, and Wang et al. 2017) or receptor mutants incapable of G protein coupling (reviewed in Aplin et al. 2009, see also Shenoy et al. 2006, Siuda et al. 2015).

Uncoupling from G protein activation was achieved for several receptors by mutating the highly conserved DRY motif found in all rhodopsin/Family A GPCRs (Wei et al. 2003, Lee et al. 2008, Bonde et al. 2010). The DRY motif, located in the cytoplasmic end of the third transmembrane helix (TM3), participates in an ionic lock with Glu in TM6 to stabilize the inactive conformation, as separation of the cytoplasmic parts of TM3 and TM6 is required for GPCR activation (Farens et al. 1996, Ballesteros et al. 2001, reviewed in Weis & Kobilka 2018). Interestingly, the prevention of this movement selectively abrogated G protein activation by the parathyroid hormone receptor (Sheikh et al. 1999, Vilardaga et al. 2001) but not GRK2-mediated receptor phosphorylation or ß-arrestin recruitment. Functional selectivity proposes that receptor can adopt multiple conformations upon ligand binding which in turn facilitate a selective activation of either G protein or ß-arrestin-dependent signaling pathways (reviewed in Smith et al. 2018, see also Littmann et al. 2015, Wang et al. 2017 and Gurevich & Gurevich 2018). In this view, charge neutralizing mutations within the DRY motif would apparently result in G protein uncoupled receptor which is, however, still capable of ß-arrestin binding and supporting ß-arrestin-mediated signaling.

In our attempt to elucidate the possibility of functional selectivity downstream of histamine 1 receptor (H1R), we engineered and characterized a H1R DRY mutant with the conserved Asp and Arg residues of the DRY motif simultaneously replaced with Ala residues. As the DRY to AAY mutation that supposedly uncouples GPCR from G protein-mediated signaling was first described for the angiotensin II type 1 receptor (AT1_A_R), we included the AT1_A_R DRY mutant in our analyses. We evaluated the subcellular localization and signaling capacity of both DRY mutants using different fluorescence resonance energy transfer (FRET)-based live cell assays. A detailed comparison of signaling dynamics downstream of the receptor was carried out. Our study sheds new light on the use of DRY mutants for studying G protein-independent signaling.

## RESULTS

### Subcellular localization of WT and DRY mutants of H1R and AT1_A_R

First we evaluated the subcellular localization of H1R DRY and AT1_A_R DRY mutants in living cells using confocal microscopy. To this end, H1R and AT1_A_R sequence were fused at their C-terminus with a red fluorescent protein, mCherry (mCh), whereas H1R DRY and AT1_A_R DRY sequence were fused similarly to a cyan fluorescent protein, mTurquoise2 (mTQ2). H1Rdry-mTQ2 and AT1_A_Rdry-mTQ2 were co-expressed with the plasma membrane marker (Lck-mVenus) and, respectively, H1R-mCh or AT1_A_R-mCh in human embryonic kidney (HEK) 293TN cells. Both H1R DRY and AT1_A_R DRY mutants showed largely overlapping subcellular localization with the respective WT receptors: they were located at the plasma membrane as well as in the intracellular (presumably endocytic) compartments. Plasma membrane localization of WT and (to a lesser extent) DRY mutant of H1R (Fig. 1A), as well as AT1_A_R and AT1_A_R DRY (Fig. 1B) was confirmed in co-localization analysis with the plasma membrane marker. The localization of WT and DRY mutant of H1R in endosomal compartment was examined by co-localization analysis with the endosomal marker, Rab7 (Vanlandingham & Ceresa 2009). To this end, H1R-mCh or H1Rdry-mCh were co-expressed with Lck-mVenus and mTQ2-Rab7 in HeLa cells. Co-localization with Rab7, indicating endosomal localization, was more pronounced in case of H1R DRY receptor than the WT H1R (Fig. 1C&D).

**Figure 1.**
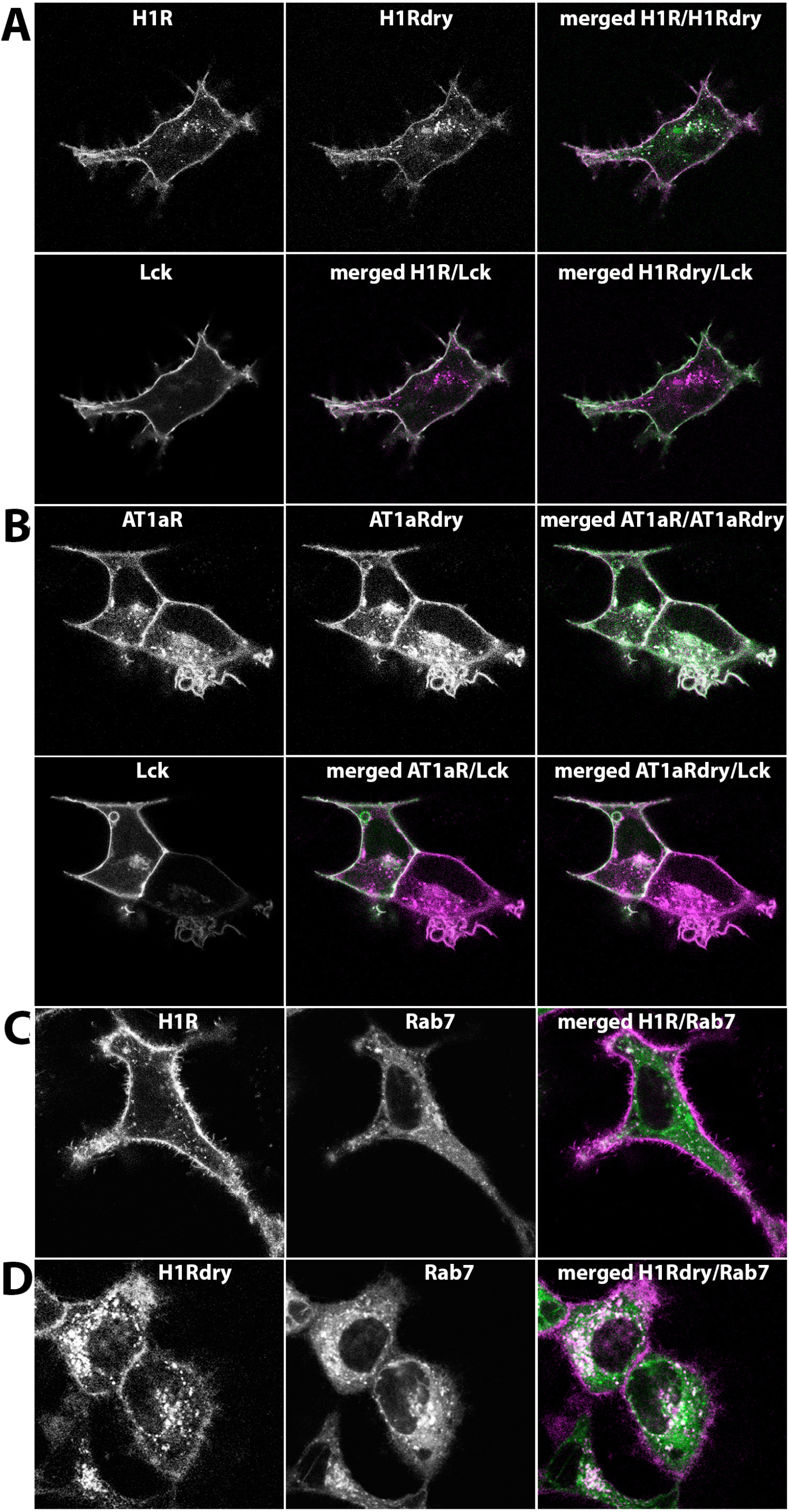
Cellular localization of WT and DRY mutants of H1R and AT1_A_R in HEK293TN cells. A) Confocal images depicting subcellular localization of WT H1R-mCh, H1Rdry-mTQ2 and Lck-mVenus co-expressed in HEK293TN cells. Upper panel (from left to right): H1R-mCh localization; H1Rdry-mTQ2 localization; merged image of H1R-mCh (set to magenta) and H1Rdry-mTQ2 (set to green) localization, where the shared localization is depicted in white. Lower panel (from left to right): localization of plasma membrane marker, Lck-mVenus; merged image of H1R-mCh (set to magenta) and Lck-mVenus (set to green) localization; merged image of H1Rdry-mTQ2 (set to magenta) and Lck-mVenus (set to green) localization. The size of the images is 60 × 60 µm. B) Confocal images depicting subcellular localization of WT AT1_**A**_R-mCh, AT1_**A**_Rdry-mTQ2 and Lck-mVenus co-expressed in HEK293TN cells. Upper panel (from left to right): AT1_**A**_R-mCh localization; AT1_**A**_Rdry-mTQ2 localization; merged image of AT1_**A**_R-mCh (set to magenta) and AT1_**A**_Rdry-mTQ2 (set to green) localization, where the shared localization is depicted in white. Lower panel (from left to right): localization of Lck-mVenus; merged image of AT1_**A**_R-mCh (set to magenta) and Lck-mVenus (set to green) localization; merged image of AT1_**A**_Rdry-mTQ2 (set to magenta) and Lck-mVenus (set to green) localization. The size of the images is 60 × 60 µm. C) Confocal images depicting subcellular localization of H1R-mCh (left) and endosomal marker, mTQ2-Rab7 (middle), co-expressed in HeLa cells. The right panel shows the merged image of H1R-mCh (set to magenta) and mTQ2-Rab7 (set to green) localization, where the shared localization is depicted in white. The size of the images is 60 × 60 µm. D) Confocal images depicting subcellular localization of H1Rdry-mCh (left) and mTQ2-Rab7 (middle) co-expressed in HeLa cells. The right panel shows the merged image of H1Rdry-mCh (set to magenta) and mTQ2-Rab7 (set to green) localization, where the shared localization is depicted in white. The size of the images is 60 × 60 µm.

### β-arrestin recruitment to WT and DRY mutants of H1R and AT1_A_R

To examine whether the DRY mutants are capable of recruiting ß-arrestins, we co-expressed WT or mutant receptors together with Lck-mVenus and ß-arr1 or ß-arr2 fused at their C-terminus with mTQ2 in HEK293TN cells. In agreement with previous reports (e.g. Oakley et al. 2000), we observed uniform cytosolic localization of both ß-arrestin fusions, with ßarr1-mTQ2, but not ßarr2-mTQ2 showing additional nuclear localization (Fig. 2). We observed rapid relocation of both isoforms to the cell periphery upon stimulation of H1R-expressing cells with 100 µM histamine, and weak ß-arrestin relocation in H1R DRY-expressing cells (Fig. 2A-D). This recruitment was visible as histamine-induced localization of ß-arrestin in discrete puncta at and near the plasma membrane (as indicated by their co-localization with Lck-mVenus) and was fully reversed upon addition of a H1R-specific antagonist, pyrilamine (PY) (see Supplemental Movies 1-4). In case of both WT and DRY mutant of AT1_A_R, stimulation with 1 µM angiotensin II (AngII) resulted in more robust and rapid ß-arr1 and ß-arr2 relocation to the plasma membrane (as indicated by their co-localization with Lck-mVenus) and subsequently to endosomal compartment (Fig. 2E-H, see Supplemental Movies 5-8), in agreement with previous reports (Oakley et al. 2000, Gaborik et al. 2003, Wei et al. 2003, Lee et al. 2008).

**Figure 2.**
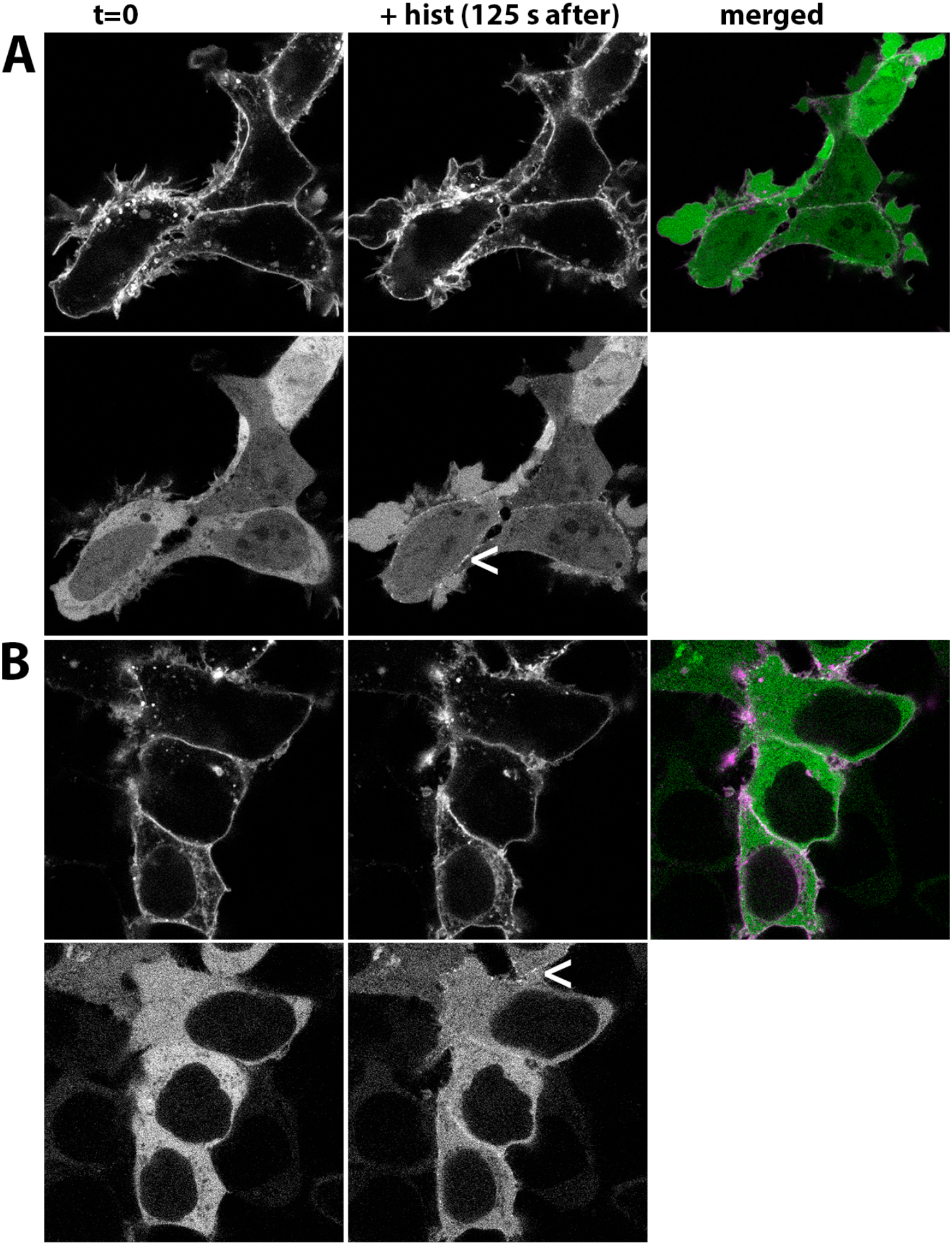
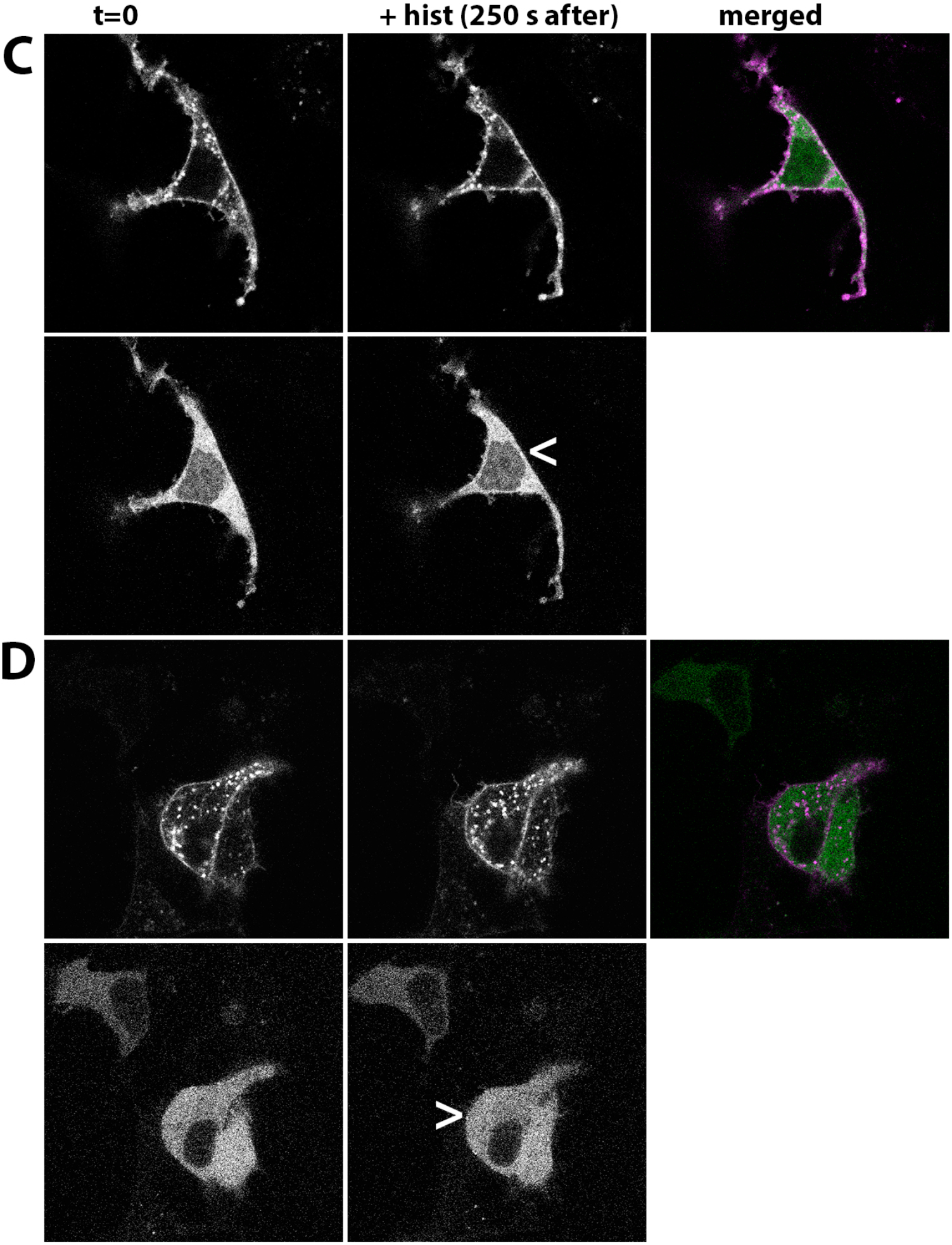
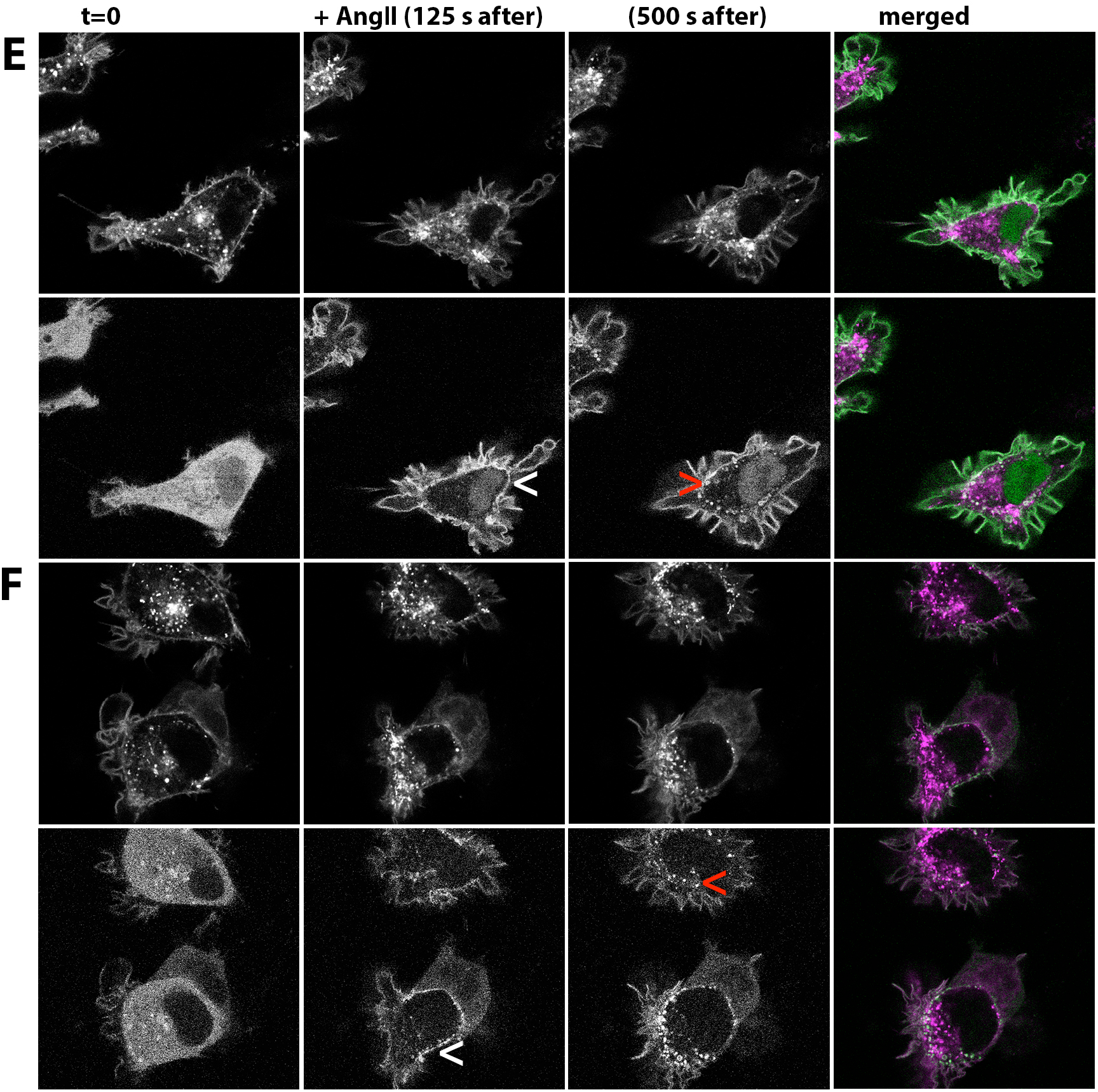
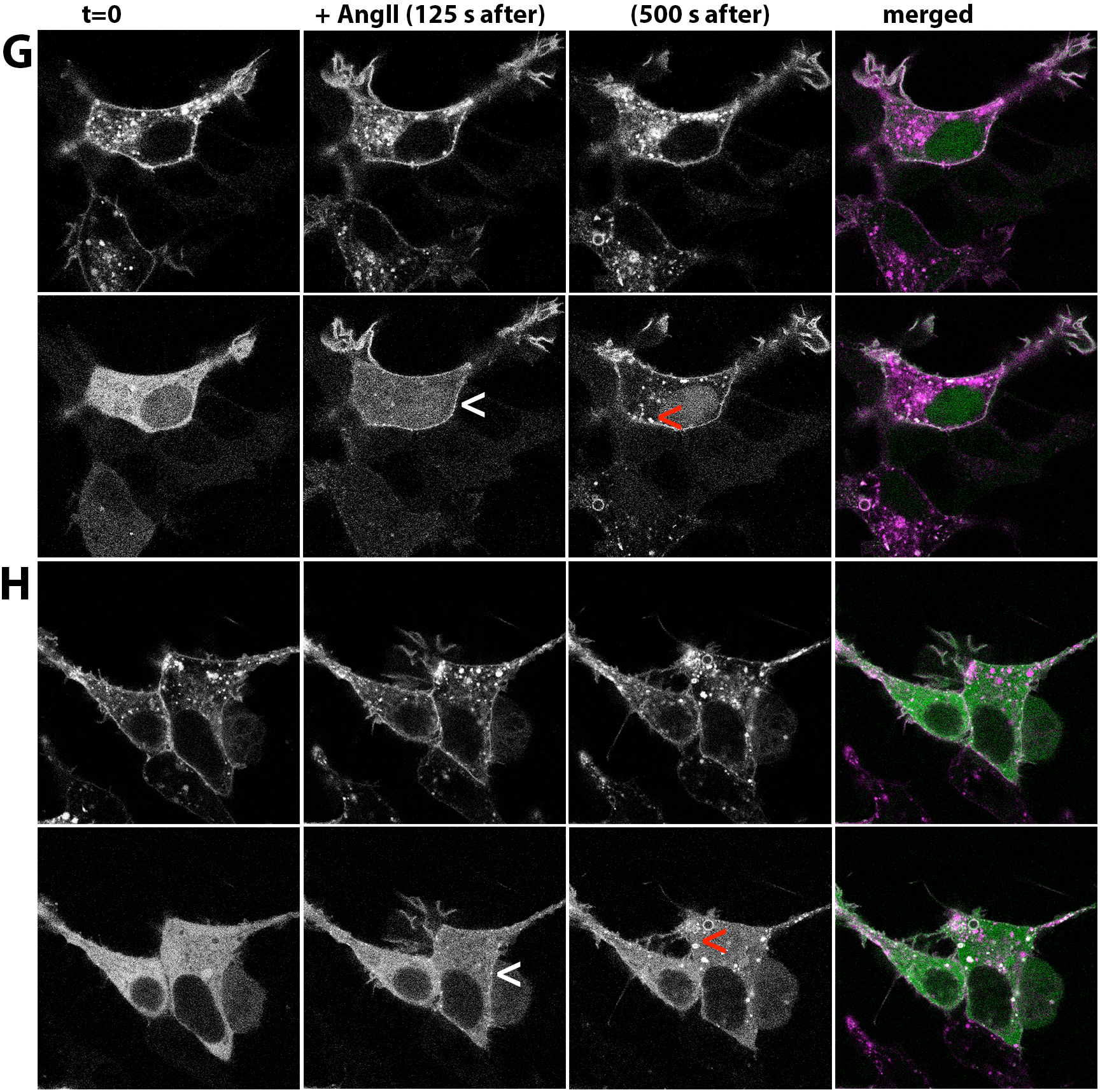
ß-arrestin recruitment upon activation of WT and DRY mutants of H1R and AT1_A_R in HEK293TN cells. A-D) Confocal images depicting subcellular localization of WT H1R-mCh (A&B, upper panels) or H1Rdry-mCh (C&D, upper panels) co-expressed together with ßarr1-mTQ2 (A&C, lower panels) or ßarr2-mTQ2 (B&D, lower panels) in HEK293TN cells. Left and middle: subcellular localization prior and after (125 s after for the WT and at 250 s after for the DRY mutant) stimulation with 100 µM histamine. Right: merged image of H1R-mCh (A&B) or H1Rdry-mCh (C&D) (set to magenta) and ßarr1-mTQ2 (A&C) or ßarr2-mTQ2 (B&D) (set to green) localization, where the shared localization is depicted in white. Arrowheads point to the histamine-induced localization of ß-arrestin at or near the plasma membrane. The size of the images is 60 × 60 µm. E-H) Confocal images depicting subcellular localization of WT AT1_**A**_R-mCh (E&F, upper panels) or AT1_**A**_Rdry-mCh (G&H, upper panels) co-expressed together with ßarr1-mTQ2 (E&G, middle panel) or ßarr2-mTQ2 (F&H, middle panel) in HEK293TN cells. From left to right: subcellular localization prior, 125 s after, and 500 s after stimulation with 1 µM AngII. Top right panel: merged image of AT1_**A**_R-mCh (E&F) or AT1_**A**_Rdry-mCh (G&H) (set to magenta) and ßarr1-mTQ2 (E&G) or ßarr2-mTQ2 (F&H) (set to green) localization 125 s (E-G, upper right panels) and 500 s (E-G, lower right panels) after AngII stimulation, where the shared localization is depicted in white. Arrowheads point to the AngII-induced localization of ß-arrestin at or near the plasma membrane (white) or in the intracellular compartment (red). The size of the images is 60 × 60 µm.

### G_q_ activation by WT and DRY mutants of H1R and AT1_A_R

Having established correct localization and the capacity to recruit β-arrestins for both the WT and the DRY mutants, we characterized functional activity of H1R DRY and AT1_A_R DRY mutants in a subset of signaling pathways. Both H1R and AT1_A_R can couple to G_q/11_ and G_i_ proteins (H1R: reviewed in Seifert et al. 2013, see also Violin et al. 2003, Adjobo-Hermans et al. 2011, van Unen et al. 2016a; AT1_A_R: reviewed in Hunyady & Catt 2006 and Kawai et al. 2017). The coupling of WT and DRY mutants to G_q_ proteins was evaluated using a FRET reporter for G_q_ activation (Adjobo-Hermans et al. 2011). Stimulation of HEK293TN cells expressing only the reporter with 100 µM histamine or 1 µM AngII did not result in any detectable FRET ratio change (Fig. 3A, see Supp. Fig. 1A for individual measurements of all cells). On the contrary, robust agonist-induced FRET signal was measured in cells co-expressing H1R-p2A-mCh (see Materials and Methods) or AT1_A_R-p2A-mCh together with the G_q_ reporter. Surprisingly, also stimulation of cells expressing H1Rdry-p2A-mCh and AT1_A_Rdry-p2A-mCh resulted in reproducible FRET signals, although they were much reduced in comparison to the responses mediated by the respective WT receptors: H1R DRY- and AT1_A_R DRY-mediated FRET signals reached, respectively, 12% and 17% of WT response. A limitation of this assay is that it requires over-expression of the G_q_ heterotrimer, i.e. Gα_q_-mTQ, Gß_1_ and YFP-Gγ_2_.

**Figure 3.**
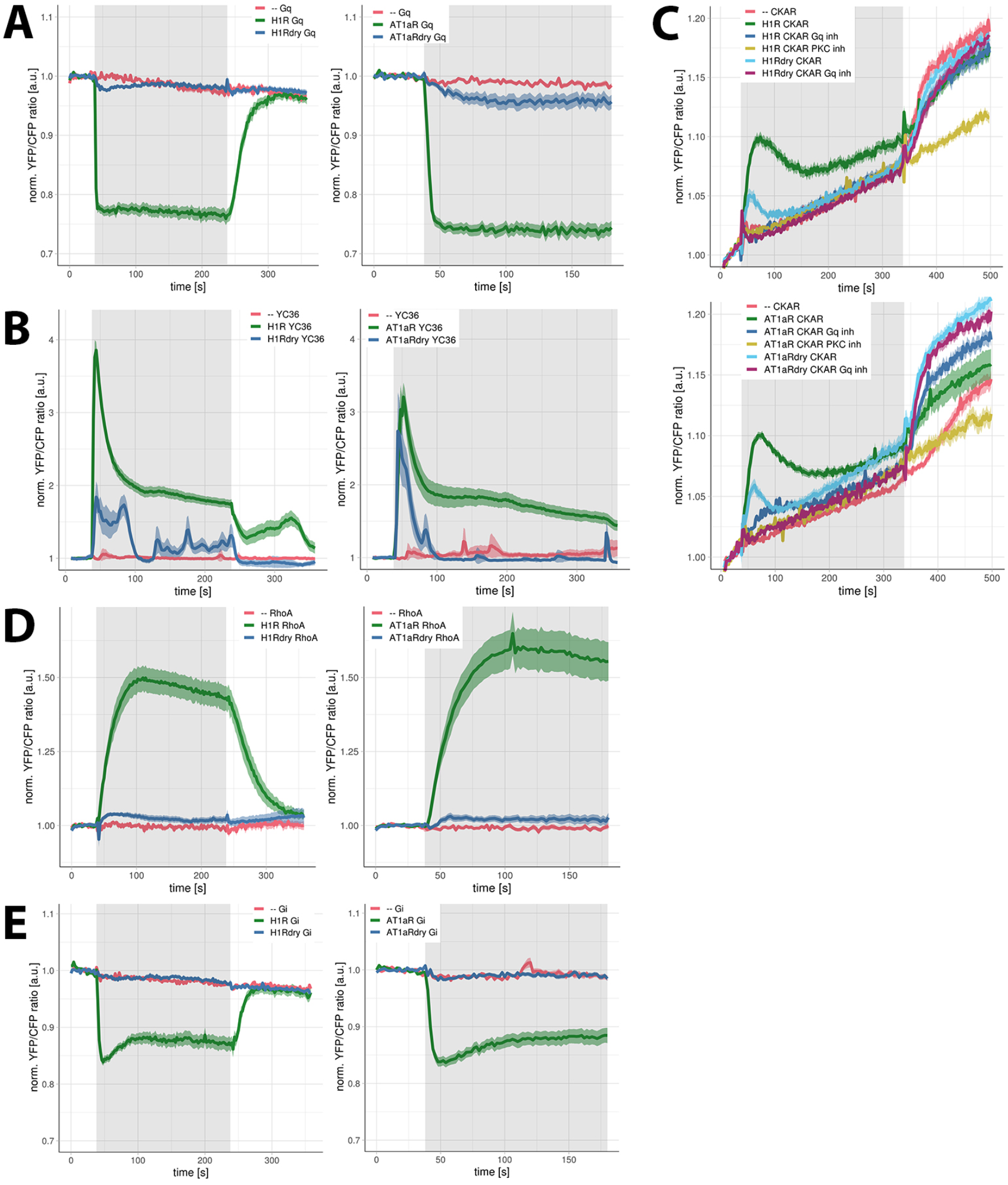
Signaling activity of WT and DRY mutants of H1R and AT1_A_R in HEK293TN cells. A) G_q_ activation by H1R, H1R DRY, AT1_**A**_R and AT1_**A**_R DRY receptors. Time traces show the average ratio change of YFP/CFP fluorescence (±95% confidence intervals) in cells expressing G_q_ sensor alone or co-expressing the sensor together with H1R-p2A-mCh, H1Rdry-p2A-mCh, AT1_**A**_R-p2A-mCh or AT1_**A**_Rdry-p2A-mCh. Left: 100 µM histamine was added at 38 s and 10 µM pyrilamine at 238 s; right: 1 µM AngII was added at 38 s. Grey boxes mark the duration of, respectively, histamine or AngII stimulation. B) Ca^2+^ changes downstream of H1R, H1R DRY, AT1_**A**_R and AT1_**A**_R DRY receptors. Time traces show the average ratio change of YFP/CFP fluorescence (±95% confidence intervals) in cells expressing YC3.6 sensor alone or co-expressing the sensor together with H1R-p2A-mCh, H1Rdry-p2A-mCh, AT1_**A**_R-p2A-mCh or AT1_**A**_Rdry-p2A-mCh. Left: 100 µM histamine was added at 38 s and 10 µM PY at 238 s; right: 1 µM AngII was added at 38 s. Grey boxes mark the duration of, respectively, histamine or AngII stimulation. C) PKC activation downstream of H1R, H1R DRY, AT1_**A**_R and AT1_**A**_R DRY receptors. Time traces show the average ratio change of YFP/CFP fluorescence (±95% confidence intervals) in cells expressing CKAR sensor alone or co-expressing the sensor together with H1R-p2A-mCh, H1Rdry-p2A-mCh, AT1_**A**_R-p2A-mCh or AT1_**A**_Rdry-p2A-mCh and untreated or treated with FR900359. 100 µM histamine (upper panel) or 1 µM AngII (lower panel) was added at 38 s and 100 nM phorbol myristate acetate (a potent PKC activator) was added at 338 s. Grey boxes mark the duration of, respectively, histamine or AngII stimulation. The observed PKC activation was abolished with a specific PKC inhibitor (10 µM Ro31-8425). Note: the continuous increase of YFP/CFP fluorescence observed in all time traces results from donor photobleaching. D) RhoA activation downstream of H1R, H1R DRY, AT1_**A**_R and AT1_**A**_R DRY receptors. Time traces show the average ratio change of YFP/CFP fluorescence (±95% confidence intervals) in cells expressing DORA-RhoA sensor alone or co-expressing the sensor together with H1R-p2A-mCh, H1Rdry-p2A-mCh, AT1_**A**_R-p2A-mCh or AT1_**A**_Rdry-p2A-mCh. Left: 100 µM histamine was added at 38 s and 10 µM PY at 238 s; right: 1 µM AngII was added at 38 s. Grey boxes mark the duration of, respectively, histamine or AngII stimulation. E) G_i1_ activation by: H1R, H1R DRY, AT1_**A**_R and AT1_**A**_R DRY receptors. Time traces show the average ratio change of YFP/CFP fluorescence (±95% confidence intervals) in cells expressing G_i_ sensor alone or co-expressing the sensor together with H1R-p2A-mCh, H1Rdry-p2A-mCh, AT1_**A**_R-p2A-mCh or AT1_**A**_Rdry-p2A-mCh. Left: 100 µM histamine was added at 38 s and 10 µM PY at 238 s; right: 1 µM AngII was added at 38 s. Grey boxes mark the duration of, respectively, histamine or AngII stimulation.

Still, our results demonstrate that the DRY mutants have guanine exchange factor activity towards a heterotrimeric G protein.

### Calcium and PKC signaling downstream of WT and DRY mutants of H1R and AT1_A_R

G_q/11_ protein-mediated activation of phospholipase Cß results in triggering of inositol triphosphate signaling and Ca^2+^-dependent protein kinase C (PKC) pathway. To evaluate G_q_ coupling of the DRY mutants in the absence of G_q_ protein over-expression, we subsequently employed FRET reporters for Ca^2+^ and PKC-dependent phosphorylation. Calcium signaling was monitored with the Yellow Chameleon 3.60 (YC3.6) FRET reporter (Fig. 3B, Supp. Fig. 1B). Stimulation of HEK293TN cells co-expressing the reporter and WT H1R or AT1_A_R with appropriate agonists resulted in immediate sharp increase of intracellular Ca^2+^ levels that subsequently decreased in a biphasic mode: the initial fast drop (to approximately 50-60% of the maximum) within 60 s of stimulation followed by much slower decrease. Stimulation of cells co-expressing the reporter and AT1_A_R DRY mutant receptors with 1 µM AngII resulted in similarly sharp increase of intracellular Ca^2+^ levels, although the response had lower amplitude and was more transient (returned to baseline within 60 s of the stimulation).

The histamine-induced YC3.6 signal in cells co-expressing the reporter and H1R DRY receptors was also much reduced in comparison to H1R-mediated response, and showed more oscillatory character (see Supp. Fig. 1B). To quantify the YC3.6 signals, we integrated all values after the addition of agonist and divided them by the number of cells measured: H1R DRY- and AT1_A_R DRY-mediated YC3.6 signals were, respectively, 21% and 17% of WT response. We measured occasional peaked Ca^2+^ increases in a subset of cells expressing only the YC3.6 sensor and stimulated with either 100 µM histamine or 1 µM AngII (Supp. Fig. 1B). These sporadic YC3.6 signals could result from the activation of endogenous H1R, H2R or AT1_A_R (presumably) expressed in HEK293TN cells (for GPCRs endogenously expressed in HEK293 cells, see Aktas et al. 2017; H2R-mediated Ca^2+^ increases were previously reported in van Unen et al. 2016a). However, the YC3.6 signals measured in cells co-expressing YC3.6 sensor and H1R DRY or AT1_A_R DRY mutant receptors were clearly different from these endogenous responses, both in kinetics and the percentage of responsive cells.

Next, the PKC activation was studied with the CKAR FRET reporter. We observed robust agonist-induced FRET ratio changes in cells co-expressing H1R or AT1_A_R together with CKAR, but not in cells expressing only the reporter (Fig. 3C, Supp. Fig. 1C). We have confirmed that the observed PKC activity was G_q_ protein-mediated, as the CKAR signal could be abolished with a Gα_q_-specific inhibitor, FR900359 (Schrage et al. 2015). CKAR signal was similarly abolished using a PKC-specific inhibitor, Ro31-8425. Using this reporter, we confirmed the ability of H1R DRY and AT1_A_R DRY to couple to G_q_ proteins: the peak of activity reached approximately 50% (for H1R DRY) or 60% (for AT1_A_R DRY) of the WT receptor-mediated response and returned faster to the baseline values (Fig. 3C).

Together, these results point to activation of G_q_ signaling by DRY mutants of the H1R and AT1_A_R, albeit at reduced levels compared to their WT counterparts.

### RhoA signaling downstream of WT and DRY mutants of H1R and AT1_A_R

As both H1 and AT1_A_ receptors can activate Rho GTPase, RhoA, via G protein (G_q/11_ for H1R, G_12/13_ for AT1_A_R)-dependent pathway (Hunyady & Catt 2006, van Unen et al. 2015), we characterized the capacity of WT and DRY mutants for RhoA activation using DORA-RhoA FRET sensor (van Unen et al. 2015). A limitation of this assay is that it requires overexpression of the RhoA. Stimulation of cells expressing only the reporter with 100 µM histamine or 1 µM AngII did not result in any detectable FRET ratio change (Fig. 3D, Supp. Fig. 1D). On the contrary, robust agonist-induced FRET signal was measured in cells co-expressing H1R or AT1_A_R together with the reporter. Stimulation of cells co-expressing the reporter and H1R DRY or AT1_A_R DRY receptors also resulted in FRET ratio changes, although these FRET signals were much weaker than those mediated by the respective WT receptors. These results are in line with the aforementioned activation of the classical G_q_ effectors, calcium and PKC, downstream of H1R DRY or AT1_A_R DRY.

### G_i_ activation by WT and DRY mutants of AT1_A_R and H1R

Additionally, we evaluated the ability of H1R DRY and AT1_A_R DRY receptors to activate G_i_ protein using a FRET sensor for G_i1_ activation (van Unen et al. 2016b). Stimulation of cells expressing only the G_i_ reporter with 100 µM histamine or 1 µM AngII did not result in any detectable FRET ratio change, whereas robust agonist-induced FRET ratio change was measured in cells co-expressing the receptor (H1R or AT1_A_R) together with the reporter (Fig. 3E, Supp. Fig. 1E). On the contrary, stimulation of cells expressing H1R DRY and AT1_A_R DRY mutant receptors did not result in detectable FRET ratio changes. Similar to the analysis using G_q_ sensor, activity of the DRY mutants on the G_i1_ protein required overexpression of the G_i_ heterotrimer.

### ERK activation by WT and DRY mutants of H1R and AT1_A_R

Finally, we characterized the ability of H1R DRY and AT1_A_R DRY receptors to activate cytosolic and nuclear pool of ERK1/2 using FRET reporters, respectively, EKARcyt and EKARnuc (Harvey et al. 2008). The localization of the reporters is shown in figure 4A. In order to measure both fast and late phases of ERK-mediated phosphorylation, we monitored EKAR signals for more than 30 minutes. Stimulation of cells expressing only EKARcyt reporter with 100 µM histamine or 1 µM AngII did not result in any FRET signal above the baseline (Fig. 4B&D, Supp. Fig. 2). In fact, we observed a transient drop (25% of maximal value) of the FRET ratio change in cells expressing only this sensor immediately after histamine stimulation. In cells expressing only the EKARnuc reporter and stimulated with 100 µM histamine or 1 µM AngII, the FRET ratio change stayed constant or showed slight increase (Fig. 4C&E, Supp. Fig. 2), rather than showing a gradual drop due to mild photobleaching (observed in vehicle-stimulated cells). This could indicate some weak agonist-induced phosphorylation of the EKARnuc reporter.

**Figure 4.**
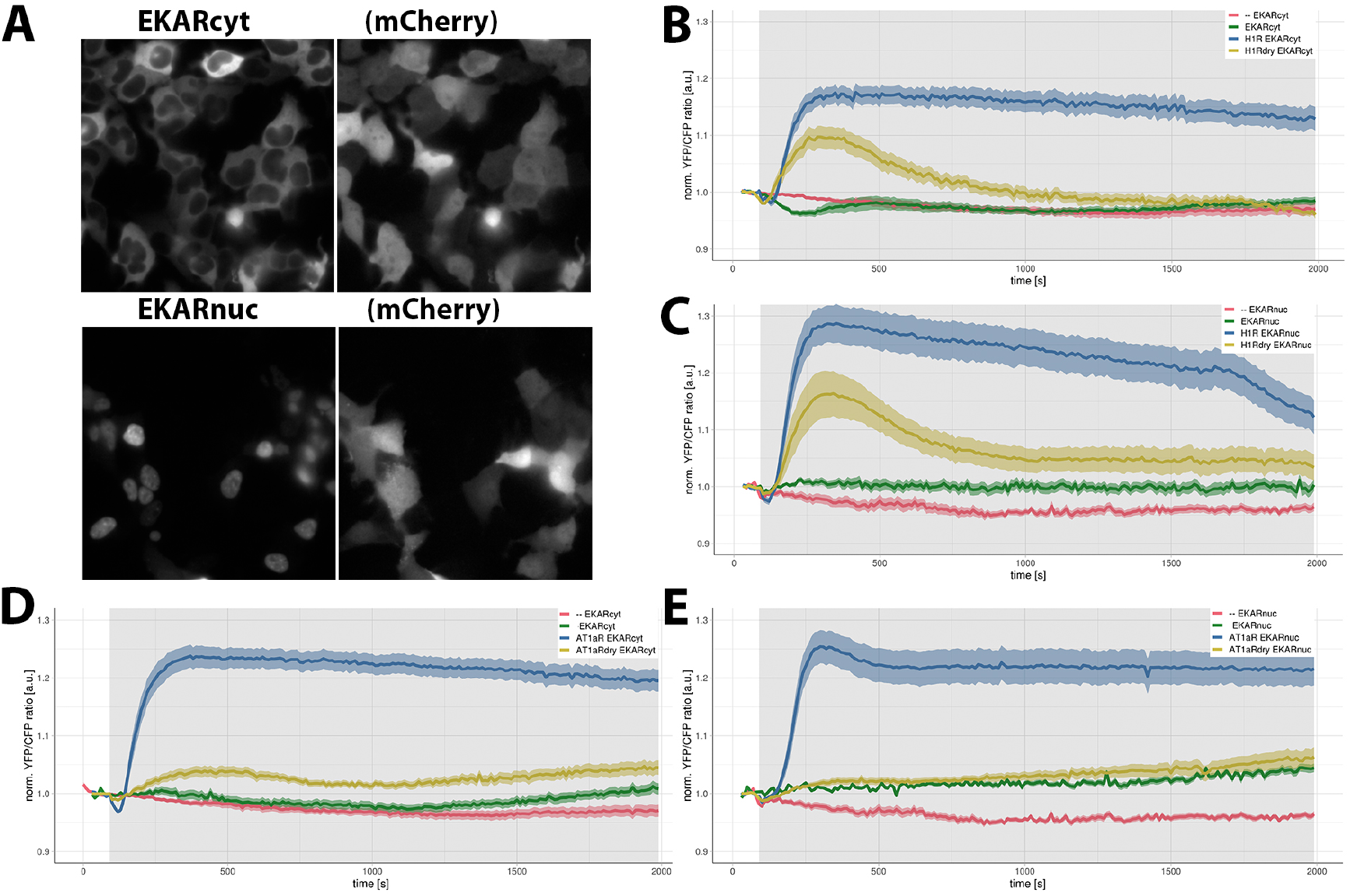
ERK1/2 activation downstream of WT and DRY mutants of H1R and AT1_A_R in HEK293TN cells. A) Wide-field images of the subcellular localization of EKARcyt and EKARnuc reporters in HEK293TN cells. Localization of the co-expressed free mCherry shows labeling of the complete cell. The size of the images is 105 x 105 µm. B&C) ERK1/2 activation downstream of WT and DRY mutant of H1R. Time traces show the average ratio change of YFP/CFP fluorescence (±95% confidence intervals) in cells expressing: EKARcyt or EKARnuc sensor and stimulated with vehicle (--) or histamine; co-expressing H1R-p2A-mCh and EKARcyt or EKARnuc and stimulated with histamine; co-expressing H1Rdry-p2A-mCh and EKARcyt or EKARnuc and stimulated with histamine. 100 µM histamine or vehicle was added at 90 s; grey boxes mark the duration of histamine stimulation. D&E) ERK1/2 activation downstream of WT and DRY mutant of AT1_**A**_R. Time traces show the average ratio change of YFP/CFP fluorescence (±95% confidence intervals) in cells expressing: EKARcyt or EKARnuc sensor and stimulated with vehicle (--) or AngII; co-expressing AT1_**A**_R-p2A-mCh and EKARcyt or EKARnuc and stimulated with AngII; co-expressing AT1_**A**_Rdry-p2A-mCh and EKARcyt or EKARnuc and stimulated with AngII. 1 µM AngII or vehicle was added at 90 s; grey boxes mark the duration of AngII stimulation.

On the contrary, we measured robust FRET signals with both EKARcyt and EKARnuc reporters in cells co-expressing the sensor together with H1R or AT1_A_R and stimulated with the respective agonist. In case of H1R-mediated ERK1/2 activation, EKARcyt and EKARnuc signals showed similar kinetics but differed in amplitude (Fig. 4B&C, Supp. Fig. 2A), with EKARnuc signal being approximately 50% higher than the EKARcyt signal. Both signals showed a transient drop immediately after histamine addition. Subsequently, both signals rose sharply, reaching maximal levels approx. 5 min after histamine addition, and then started to slowly decay. H1R DRY-mediated ERK activation, although reduced, resembled the response downstream of WT H1R (Fig. 4B&C, Supp. Fig. 2A): both EKARcyt and EKARnuc signals showed initial drop, after which they started increasing, reaching more than 50% of the respective H1R-mediated responses within 5 min of histamine addition. H1R DRY-mediated EKAR signals showed however much faster decline than the H1R-mediated signals.

Stimulation of cells expressing EKARcyt or EKARnuc reporter and AT1_A_R with 1 µM AngII (Fig. 4D&E; Supp. Fig. 2B) resulted in EKAR signals resembling histamine-induced responses of H1R-expressing cells. However, AngII-induced responses differed in amplitude from the respective histamine-induced responses (AngII-induced EKARcyt signal being higher and EKARnuc signal lower), and showed slower decay. Both EKARcyt and EKARnuc signals showed similar kinetics and amplitude, although a transient drop (13% of maximal value) immediately upon AngII addition was observed only with the former reporter. Interestingly, cells expressing AT1_A_R DRY and stimulated with AngII were only able to activate the cytosolic ERK pool (Fig. 4D&E, Supp. Fig. 2B). We did not measure any AT1_A_R DRY-mediated EKARnuc signal above the response of cells expressing only EKARnuc reporter and stimulated with AngII.

## DISCUSSION

Engineered GPCRs that have a mutation in the DRY motif have been proposed as tools to study G-protein independent signaling. It is important to verify to what extent mutations in the DRY motif inhibit G-protein signaling. Here, we report the signaling dynamics of two class A receptors, H1R and AT1_A_R, in which the DRY motif is changed to AAY. Our results demonstrate that DRY mutants of H1R and AT1_A_R are capable of activating heterotrimeric G-proteins, resulting in downstream signaling, including ERK activation.

The DRY motif in AT1_A_R is required for receptor activation, although there are contradictory results regarding the level of impairment achieved with identical or similar mutations. Ohyama and colleagues (2002) using Chinese hamster ovary (CHO-K1) cells reported severe reduction in G protein coupling (as indicated by the insensivity of AngII binding to GTPγS) and inositol phosphate (IP) production for AT1_A_R mutants with either Asp125 or Arg126 residue replaced by either Ala or Gly. On the contrary, Gaborik and colleagues (2003) using COS-7 cells observed severe impairment of IP production and ERK1/2-dependent Elk1 promoter activation only with double DRY/AAY mutant. Importantly, AT1_A_R DRY/AAY or DRY/GGY double mutants expressed in HEK293 or COS-7 cells were reported to be unable to activate G protein (measured at the level of ^35S^GTPγS binding, IP production or Ca^2+^ accumulation) but capable of ß-arrestin recruitment and ERK1/2 activation (Wei et al. 2003, Lee et al. 2008, Bonde et al. 2010). However, residual activity (approx. 25% of the WT response measured at the maximal value) of DRY/AAY mutant in FLIPR assay that measures increases of intracellular Ca^2+^ levels was observed (Lee et al. 2008). Additionally, Bonde and colleagues (2010) reported increased Sar^1^-Ile^4^-Ile^8^ (SII) AngII-induced IP production downstream of DRY/AAY mutant activation compared to WT AT1_A_R in COS-7 cells, indicating the ability of this receptor mutant to couple to G_q/11_ protein.

Here, we observed substantial, albeit reduced compared to WT receptor responses, G_q_ protein coupling of H1R DRY and AT1_A_R DRY mutants in HEK293TN cells using G_q_ FRET reporter. The fact that we did not observe similar coupling of these mutants to G_i1_ could suggests signaling bias (preference for G_q_ coupling over G_i1_). However, it could also result from lower dynamic range of the G_i1_ FRET reporter. The observed H1R DRY and AT1_A_R DRY coupling to G_q_ protein was achieved in cells transiently expressing all three subunits of the G_q_ heterotrimer. Previously, overexpression of Gα_q_ protein was shown to increase AngII-induced signaling for all tested AT1_A_R mutants except the DRY/AAY mutant (Bonde et al. 2010). To confirm coupling of these receptor mutants to G_q_ protein in the absence of its overexpression, we characterized signaling ability of H1R DRY and AT1_A_R DRY receptors using FRET reporters for G_q_ effectors, i.e. PKC and Ca^2+^. Both CKAR and YC3.6 sensors reported efficient coupling of H1R DRY and AT1_A_R DRY mutants to G_q_ protein. Signaling efficiencies of these mutant receptors measured with CKAR and YC3.6 sensors were increased in comparison to G_q_ FRET signal, likely as a result of amplification of the signaling outcome downstream of the initial (reduced) coupling of these receptor mutants to G_q_ proteins.

H1R DRY and AT1_A_R DRY receptors were also able to weakly activate RhoA. Endogenous H1 receptors in HeLa cells were shown to activate RhoA through p63RhoGEF- or Trio-mediated, G_q_ protein-dependent mechanism (van Unen et al. 2015). Similarly, AT1_A_R was shown to increase RhoA activity via the G_12/13_-dependent, RhoGEF-mediated pathway in vascular smooth muscle cells and in cardiac myocytes in culture and in vivo (reviewed in Kimura and Eguchi 2009). However, Bregeon and colleagues (2009) have recently demonstrated G protein-independent activation of RhoA downstream of AT1_A_R in cultured vascular smooth muscle cells. As we have not investigated the ability of H1R DRY and AT1_A_R DRY mutants to activate RhoA in the presence of G_q_ or G_12/13_ protein inhibitors (FR900359 and RGS domain of p115RhoGEF, respectively), we cannot currently conclude on the mechanism of this activation.

Taken together, our results demonstrate the ability of H1R DRY and AT1_A_R DRY mutant receptors to signal via G_q_ proteins, with the strength of the signal strongly depending on the signaling event tested (with more downstream effectors showing higher signals due to the signal amplification). At the same time, we have noted a very transient character of this G_q_ coupling of the DRY mutants: histamine-induced G_q_ and CKAR signals, as well as AngII-induced CKAR and YC3.6 signals decayed within a minute of agonist stimulation. Such transient signals would likely be missed or blunted in conventional biochemical assays that generally have a poor temporal resolution. Therefore, our results once again demonstrate the strength of single cell imaging using biosensors (Greenwald et al. 2018).

With respect to the ß-arrestin recruitment, we observed agonist-induced relocation of both ß-arrestin isoforms to cell periphery of H1R DRY- and AT1_A_R DRY-expressing HEK293TN cells. H1R DRY-mediated relocation was however less robust than ß-arrestin recruitment upon WT H1R activation. Previously, bioluminescence resonance energy transfer between ßarr2-GFP (green fluorescent protein) fusion protein and AT1_A_R DRY fusion to Renilla luciferase, indicating receptor-ßarr2 interaction (or at least close proximity), was reported by Bonde and colleagues (2010). Agonist-induced relocation of ßarr1− and/or ßarr2-GFP fusion proteins in AT1_A_R- and AT1_A_R DRY-expressing cells was also demonstrated using confocal microscopy (Wei et al. 2003, Lee et al. 2008). Both research groups showed the change in ß-arrestin cellular distribution, from uniformly cytosolic to presumably sequestered in endocytotic compartment, upon agonist stimulation. However, this redistribution was only demonstrated after 30 minutes of stimulation and in fixed cells. On the contrary, our results showed agonist-induced relocation of both ß-arrestin isoforms to the cell periphery as early as 2 minutes of stimulation. Additionally, we reported overlap of ß-arrestin puncta with the receptor location at the cell periphery and, in case of AT1_A_R- and AT1_A_R DRY-expressing cells, also in intracellular (presumably endosomal) compartment, indicating co-localization of ß-arrestin and receptor fluorescent fusions.

Finally, we confirmed the ability of WT and DRY mutant receptors to activate ERK. Importantly, AngII-induced EKAR signals (both EKARcyt and EKARnuc), reporting on ERK-mediated phosphorylation, significantly resembled AngII-induced ERK phosphorylation demonstrated in HEK293 cells stably expressing AT1_A_R using conventional immunoblotting (Ahn et al. 2004). We similarly observed an immediate increase of phosphorylation upon 1 µM AngII addition, reaching maximum value at 5 minutes of stimulation, and followed by a slow decay. The observed drop of AT1_A_R-mediated EKAR signals achieved at 30 minutes of stimulation was lower than the 25% decrease of ERK phosphorylation reported in Ahn et al. 2004 (EKARcyt and EKARnuc signals reached at that time 83% and 84% of maximal response, respectively). H1R-mediated EKARcyt and EKARnuc signals reached, respectively, 76% and 53% of the maximal response at 30 min of stimulation. Therefore, our results support the notion that EKAR sensors can faithfully report ERK activation in living cells. An intriguing observation was the transient drop in histamine-induced EKARcyt and EKARnuc signals, as well as AngII-induced EKARcyt signal, indicating agonist-induced transient inhibition of ERK activity or dephosphorylation of the existing phosphoERK pool.

The G protein-dependent and ßarrestin-dependent ERK activation was postulated to differ both with respect to its kinetics and spatial distribution (Luttrell et al. 2001, Tohgo et al. 2002, Ahn et al. 2004, Ren et al. 2005, Shenoy et al. 2006). In HEK293 cells expressing AT1_A_R, beta2 adrenergic receptor or vasopressin V2 receptor, G protein-dependent activation was reported to be rapid in onset but transient (with most of the signal waning within 10 min of stimulation), and to generate phosphorylated ERK distributed throughout the cytosol and nucleus. On the contrary, ßarrestin-dependent component was reported to reach maximum at 10 min, persist for at least 30 min without attenuation, and activate only the cytosolic ERK1/2 pool.

We measured reproducible agonist-induced EKARcyt signals downstream of both H1R DRY and AT1_A_R DRY. AT1_A_R DRY-mediated EKARcyt signal (16% of the maximal value) was much lower than the AT1_A_R DRY-mediated ERK phosphorylation measured in HEK293 cells using conventional methods (25% in Lee et al. 2008, 50% in Wei et al. 2003, 75% in Ahn et al. 2004) but similarly persistent (Ahn et al. 2004, Lee et al. 2008). Currently, we are not able to conclude on the reason of the observed discrepancy between our results and those obtained by other research groups. With respect to ERK activation in the nucleus, we did not observe any agonist-induced EKARnuc signal in AT1_A_R DRY-expressing cells, in agreement with the lack of ERK-mediated gene expression downstream of AT1_A_R DRY stimulation reported previously (Gaborik et al. 2003, Lee et al. 2008). On the contrary, histamine-induced EKARnuc signal in cells expressing H1R DRY mutant receptor achieved 60% of the WT response. It would be interesting to find out if this nuclear ERK activity also results in histamine-induced transcriptional reprogramming.

Several research groups postulated the existence of a G protein-independent, ß-arrestin-dependent ERK activation downstream of several GPCRs, including AT1_A_R (reviewed in Aplin et al. 2009 and Whalen et al. 2011, see also Wei et al. 2003, Shenoy et al. 2006, Violin et al. 2010, Zimmerman et al. 2012; for the apparent capacity of “ß-arrestin biased” agonist of AT1_A_R to activate G protein-mediated signaling, see Saulière et al. 2012). Some of these studies employed the DRY receptor mutants to claim G protein independence of this ERK activation. However, residual ability of AT1_A_R DRY and H1R DRY mutant receptors demonstrated in this study challenges the notion that the DRY/AAY mutation leads to complete uncoupling of the receptor from G proteins. Importantly, stimulation of different GPCRs in the absence of G protein activation (including “zero functional G” background) was recently reported not to result in any ERK activation (Alvarez-Curto et al. 2016, Grundmann et al. 2018), also challenging the concept of G protein-independent ERK activation. It would be interesting to investigate whether distinct ERK activation observed downstream of DRY mutant activation could be reproduced by mimicking suboptimal and transient activation of G_q_ protein, such as demonstrated in this study.

## MATERIALS and METHODS

### Constructs

Plasmids encoding HsH1R-mCherry and H1R-p2A-mCherry are described in van Unen et al. (2016a). The p2A viral sequence in the latter construct ensures that the mCherry is separated from the receptor protein during translation, generating untagged receptor and free RFP that reports on receptor translation levels. Plasmid encoding RnAT1_A_R-mVenus was a kind gift from Peter Várnai (Semmelweis University, Hungary). The AT1_A_R sequence was cloned into pN1-mCherry and pN1-p2A-mCherry vectors using HindIII and AgeI sites. The DRY/AAY mutation was introduced into H1R and AT1_A_R sequence using site-directed mutagenesis; after the mutations were confirmed with sequencing, H1R DRY and AT1_A_R DRY sequence were cloned into pN1-mCherry and pN1-p2A-mCherry vectors (Clontech). Plasmids encoding Rnßarr1-mYFP (plasmid #36916) and Rnßarr2-mYFP (plasmid #36917) were purchased from addgene.org and. Plasmids encoding ßarr1-mTQ2 and ßarr2-mTQ2 were generated by cloning ß-arr1 and ß-arr2 sequences into pN1-mTurquoise2 Clontech vector using PCR amplification and KpnI and AgeI sites. Plasmids encoding all H1R, AT1 _A_R, ß-arr1 and ß-arr2 constructs used in this study will be deposited at http://www.addgene.org. Plasmids encoding the Gq sensor (Goedhart et al. 2011) and the DORA-RhoA sensor (Unen et al., 2015) were reported before. Plasmids encoding the endosomal marker, mTurquoise2-Rab7, the calcium biosensor YC3.6, and the G_i1_ biosensor are available from http://www.addgene.org. Lck-mVenus was reported as a plasma membrane marker in van Unen et al. (2016b). CKAR biosensor (Violin et al. 2003) was a kind gift from Alexandra Newton (University of California, San Diego). Cerulean-Venus versions of the EKARcyt (#18679) and EKARnuc (#18681) biosensors (Harvey et al. 2008) were purchased from addgene.org.

### Reagents

Histamine, angiotensin II (AngII), pyrilamine (PY), phorbol myristate acetate (PMA), pertussis toxin (PTX), and Ro31-8425 were purchased from Sigma Aldrich/Merck. The specific Gαq inhibitor, FR900359 (UBO-QIC), was purchased from the University of Bonn (http://www.pharmbio.uni-bonn.de/signaltransduktion). FR900359 was added to cells (in microscopy medium) at least 10 minutes before the measurements at a concentration of 1 µM. Ro31-8425 was added to cells (in microscopy medium) at least 25 minutes before the measurements at a concentration of 10 µM. Experiments with PTX inhibitor were carried out using cells cultured in serum free DMEM and incubated o/n with 100 ng/µL PTX.

### Cell culture & sample preparation

HEK293TN cells (System Biosciences, LV900A-1) and HeLA cells (American Tissue Culture Collection: Manassas, VA, USA) were cultured using Dulbecco’s Modified Eagle Medium (DMEM) supplied with glutamax, 10% FBS, penicillin (100 U/ml) and streptomycin (100 ug/ml), and incubated at 37°C and 5% CO_2_. All cell culture regents were obtained from Invitrogen (Bleiswijk, NL). Cells were transfected in a 35 mm dish holding a glass coverslip (24 mm Ø, Menzel-Gläser, Braunschweig, Germany), using polyethylenimine (3 μL of PEI:1 μL of DNA) according to the manufacturer’s protocol. For each transfection, we used 500 ng of receptor (H1R, H1R DRY, AT1_A_R or AT1_A_R DRY)-carrying plasmid. Other plasmids were transfected at: 100 ng (ßarr1 fusion constructs), 150 ng (ßarr2 fusion constructs), 200 ng (Lck-mVenus, CKAR, YC3.6), 250 ng (EKARcyt, EKARnuc), 300 ng (DORA-RhoA), 750 ng (G_q_ reporter, G_i1_ reporter). Samples were imaged one day after transfection: coverslips were mounted in an Attofluor cell chamber (Invitrogen, Breda, NL) and submerged in 1 mL microscopy medium (20 mM HEPES, pH=7.4, 137 mM NaCl, 5.4 mL KCl, 1.8 mM CaCl_2_, 0.8 mM MgCl_2_, and 20 mM glucose; Sigma Aldrich/Merck).

### Confocal and wide-field microscopy

Relocation experiments were performed as described in van Unen et al. (2016c). Ratiometric FRET measurements were performed on a previously described wide-field microscope (van Unen et al. 2015). Typical exposure time was 100 ms, and camera binning was set to 4×4. Fluorophores were excited with 420/30 nm light and reflected onto the sample by a 455DCLP dichroic mirror. CFP emission was detected with a BP470/30 filter and YFP emission was detected with a BP535/30 filter by rotating the filter wheel. RFP was excited with 570/10 nm light reflected onto the sample by a 585 dichroic mirror, and RFP emission was detected with a BP620/60 nm emission filter. All acquisitions were corrected for background signal and bleedthrough of CFP emission in the YFP channel. All experiments were performed at 37°C.

### Image analysis and data visualization

ImageJ (National Institute of Health) was used to analyze the raw microscopy images. Background subtractions, bleedthrough correction and calculation of the normalized ratio per time point per cell were done in Excel (Microsoft Office). All plots were prepared with the PlotTwist web app (Goedhart 2019). Plots show the average response as a thicker line and a ribbon for the 95% confidence interval around the mean.

## Author contributions

APB designed and performed experiments, analyzed the data, and wrote the manuscript. LJ provided technical assistance with cell culture maintenance and confocal microscopy. JG participated in study design, data interpretation, and data visualization. All authors approved the final manuscript.

## ACKNOWLEDGEMENTS

We thank Peter Várnai (Semmelweis University, Hungary) and Alexandra Newton (University of California, San Diego) for providing constructs.

## SUPPLEMENTARY DATA

**Supplemental Figure 1:**
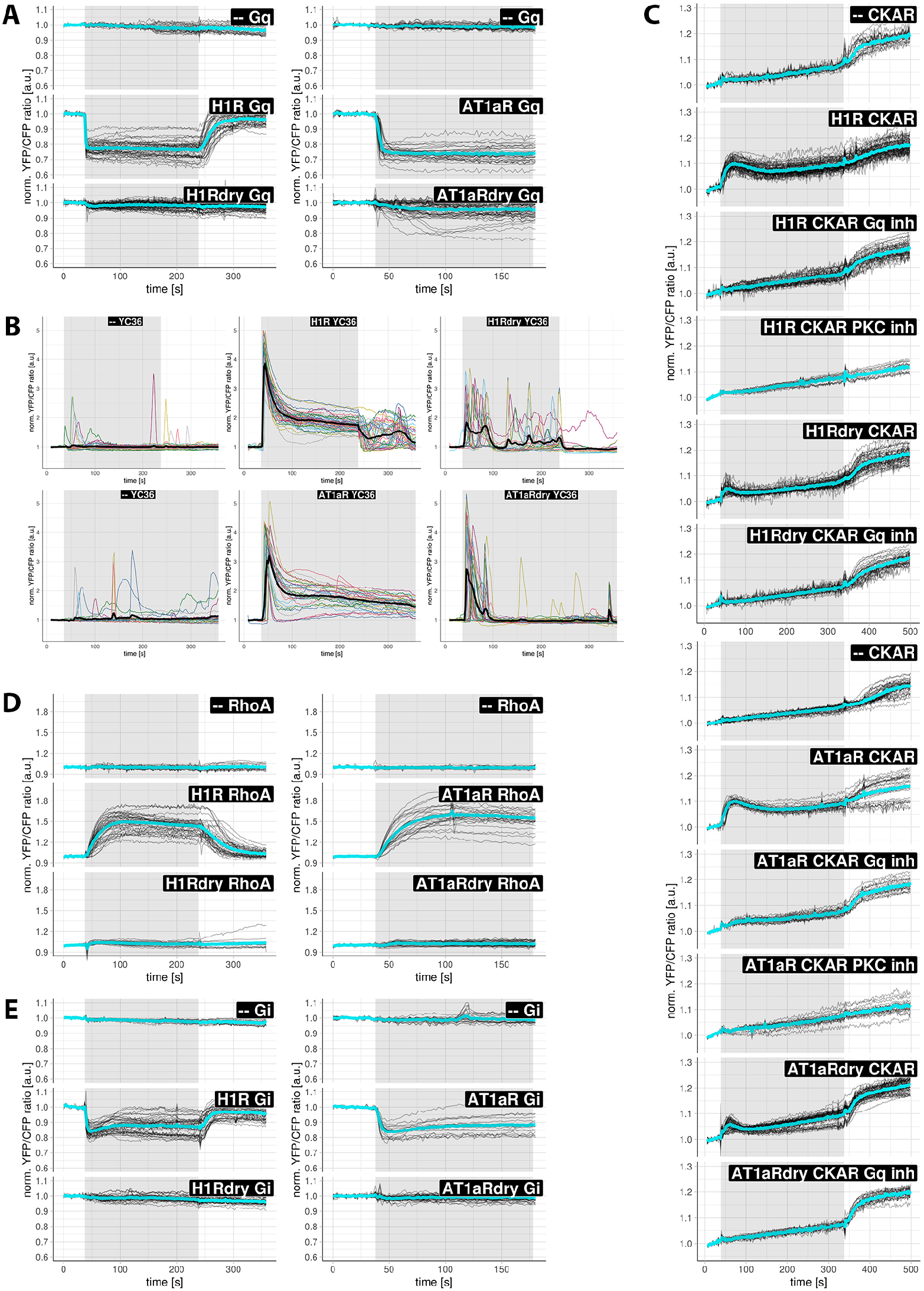
Signaling activity of WT and DRY mutants of H1R and AT1_A_R in HEK293TN cells. Black (A, C-E) or color (B) lines show the ratio change of YFP/CFP fluorescence in individual cells, and the average ratio change is shown as a thicker cyan (A, C-E) or black (B) line. Gq (A), YC3.6 (B), CKAR (C), DORA-RhoA (D) or Gi (E) sensor was expressed alone (--) or co-expressed together with H1R-p2A-mCh, H1Rdry-p2A-mCh, AT1_**A**_R-p2A-mCh or AT1_**A**_Rdry-p2A-mCh; black boxes specify the constructs (co-)expressed in cells. C) Cells were untreated, treated with a specific Gq inhibitor (FR900359) or with a specific PKC inhibitor (Ro31-8425). (A&B and D&E, left) 100 µM histamine was added at 38 s and 10 µM PY at 238 s, except in C (top and bottom), where 100 nM phorbol myristate acetate (PMA) was added at 338 s; (A&B and D&E, right) 1 µM AngII was added at 38 s. Grey boxes mark the duration of, respectively, histamine or AngII stimulation.

**Supplemental Figure 2:**
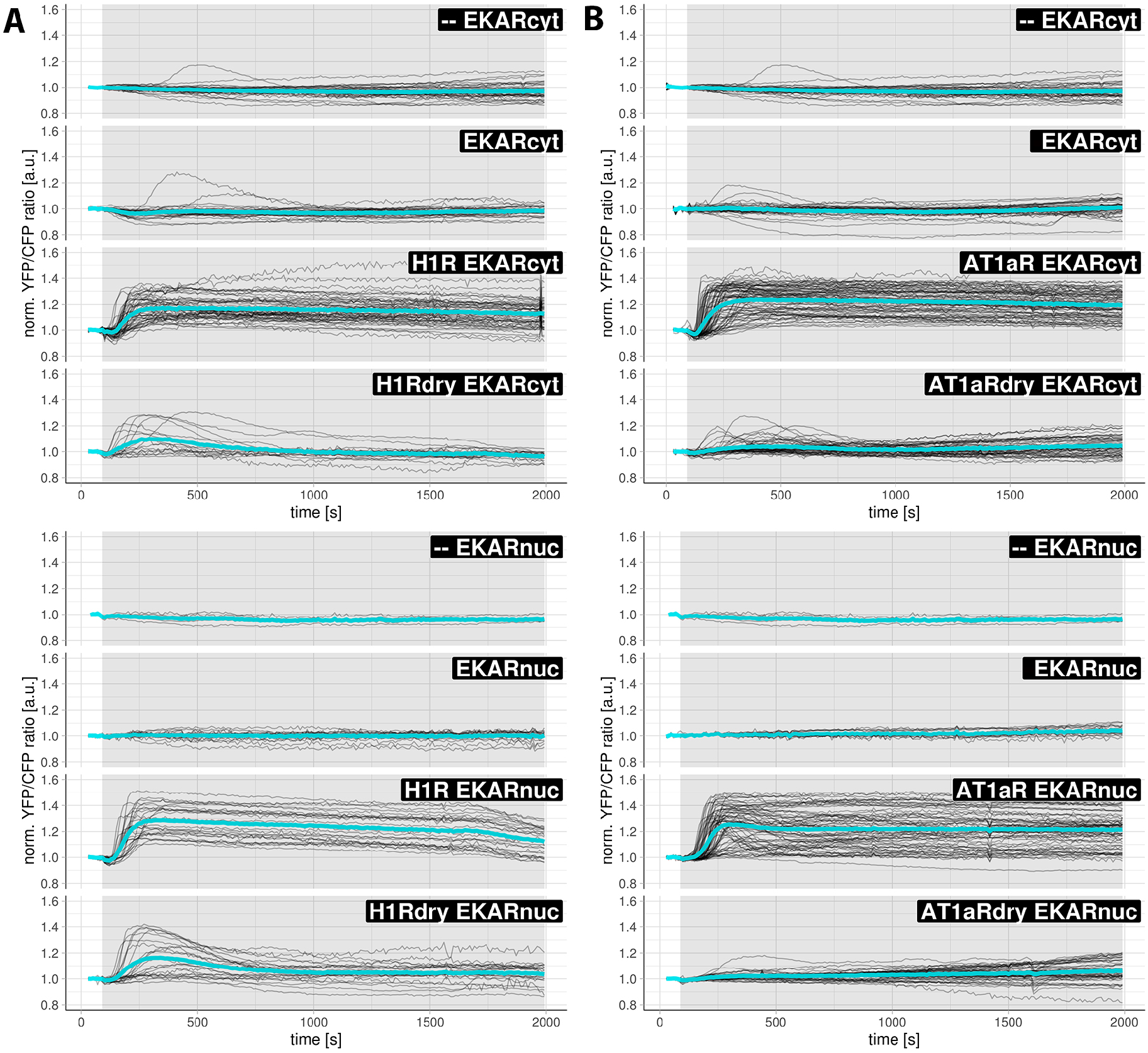
ERK activation downstream of WT and DRY mutants of H1R and AT1_A_R in HEK293TN cells. Black lines show the ratio change of YFP/CFP fluorescence in individual cells, and the average ratio change is shown as a thicker cyan line. EKARcyt (A&B, upper panels) or EKARnuc (A&B, lower panels) were expressed alone or co-expressed together with H1R-p2A-mCh, H1Rdry-p2A-mCh, AT1_**A**_R-p2A-mCh or AT1_**A**_Rdry-p2A-mCh; black boxes specify the constructs (co-)expressed in cells. Vehicle (--), 100 µM histamine (A) or 1 µM AngII (B) was added at 90 s; grey boxes mark the duration of histamine or AngII stimulation, respectively.

**Supplemental Movies. ß-arrestin recruitment to plasma membrane upon H1R or H1R DRY activation in HEK293TN cells.**

Movie 1-4) Confocal recordings of subcellular localization of ßarr1-mTQ2 (1&3) or ßarr2-mTQ2 (2&4) co-expressed together with H1R-mCh (1&2) or H1Rdry-mCh (3&4) in HEK293TN cells. 100 µM histamine was added at 225 s and 10 µM PY at 975 s. Time interval is 25 s. The size of the images is 60 × 60 µm.

Movie 5-8) Confocal recordings of subcellular localization of ßarr1-mTQ2 (5&7) or ßarr2-mTQ2 (6&8) co-expressed together with AT1_**A**_R-mCh (5&6) or AT1_**A**_Rdry-mCh (7&8) in HEK293TN cells. 1 µM AngII was added at 225 s. Time interval is 25 s. The size of the images is 60 × 60 µm.

